# Infant antibody repertoires during the first two years of influenza vaccination

**DOI:** 10.1101/2022.09.13.507806

**Authors:** Masayuki Kuraoka, Nicholas C. Curtis, Akiko Watanabe, Hidetaka Tanno, Seungmin Shin, Kevin Ye, Elizabeth Macdonald, Olivia Lavidor, Susan Kong, Tarra Von Holle, Ian Windsor, Gregory C. Ippolito, George Georgiou, Emmanuel B. Walter, Garnett H. Kelsoe, Stephen C. Harrison, M. Anthony Moody, Goran Bajic, Jiwon Lee

**Affiliations:** Department of Immunology, Duke University, Durham, NC 27710, USA; Thayer School of Engineering, Dartmouth College, Hannover, NH 03755, USA; Departments of Chemical Engineering, Molecular Biosciences, and Biomedical Engineering, and Institute for Cellular and Molecular Biology, University of Texas, Austin, TX 78712, USA; Laboratory of Molecular Medicine, Boston Children’s Hospital and Harvard Medical School, Boston, MA 02115, USA; Department of Pediatrics and Duke Human Vaccine Institute, Duke University, Durham, NC 27710, USA; Department of Molecular Biosciences and Institute for Cellular and Molecular Biology, University of Texas, Austin, TX 78712, USA; Department of Microbiology, Icahn School of Medicine at Mount Sinai, New York, NY 10029, USA; Howard Hughes Medical Institute, Harvard Medical School, Boston, MA 02115, USA

## Abstract

The first encounter with influenza virus biases later immune responses. This “immune imprinting”, formerly from infection within a few years of birth, is in the U.S. now largely from immunization with a quadrivalent, split vaccine (IIV4). In a pilot study of IIV4 imprinting, we characterized, by single-B-cell cultures, NextGen sequencing, and plasma antibody proteomics, the primary antibody responses to influenza in two infants during their first two years of seasonal influenza vaccination. One infant, who received only a single vaccination in Year 1, contracted an influenza B (IBV) infection between the two years, allowing us to compare imprinting by infection and vaccination. That infant had a shift in hemagglutinin (HA)-reactive B-cell specificity from largely influenza A (IAV)-specific in Year 1 to IBV-specific in Year 2, both before and after vaccination. HA-reactive B cells from the other infant maintained a more evenly distributed specificity. In Year 2, class-switched HA-specific B cell *IGHV* somatic hypermutation (SHM) levels reached average levels seen in adults. The HA-reactive plasma antibody repertoires of both infants comprised a relatively small number of antibody clonotypes, with one or two very abundant clonotypes. Thus, after the Year 2 boost, both infants had overall B cell profiles that resembled those of adult controls.

**Importance:** Influenza virus is a moving target for the immune system. Variants emerge that escape protection from antibodies elicited by a previously circulating variant (“antigenic drift”). The immune system usually responds to a drifted influenza virus by mutating existing antibodies rather than by producting entirely new ones. Thus, immune memory of the earliest influenza exposure has a major influence on later responses to infection or vaccination (“immune imprinting”). In the many studies of influenza immunity in adult subjects, imprinting has been from an early infection, since only in the past two decades have infants received influenza immunizations. The work reported in this paper is a pilot study of imprinting in two infants by the flu vaccine, which they received before experiencing an influenza infection. The results suggest that a quadrivalent (four-subtype) vaccine may provide an immune imprint less dominated by one subtype than does a monovalent infection.

## Introduction

Influenza A (IAV) and B (IBV) viruses that circulate in the human population evolve in response to acquired global host immunity. Antigenic drift, from mutation and selection for resistance against existing immunity, generates a new “antigenic cluster” every 5 to 10 years, depending on the influenza type and subtype (1). Most of such variation occurs within the hemagglutinin (HA) protein.

Classic epidemiological and serological studies, as well as contemporary molecular analyses, indicate that the antigenic variant of an influenza virus subtype first encountered conditions later responses -- a phenomenon called “original antigenic sin” in early papers and now known as “immune imprinting” (2-5). The presumed mechanism is that immune memory from the earliest exposure, both in the form of memory B cells (Bmem) and long-lived plasma cells (LLPCs, which secrete and maintain steady-state plasma antibodies), governs the immune outcome of all subsequent exposures. In accord with such a mechanism, lineage reconstructions have shown that B cells elicited by vaccination of adults are descendants of those elicited by the presumptive infecting strain circulating at the time of the donors’ birth (6). Less clear are the extent to which the subtype of the initial exposure dominates the response to multivalent vaccines and whether the B cell memory established by exposure to a new subtype (*e*.*g*., H3 in 1968 for those imprinted earlier by H1) is primarily a naive response or rather modification or adaptation of Bmem from the initial imprinting. The usually uncertain history of various exposures to IAV and IBV between early childhood and sample collection has confounded interpretation of most data from individual adult donors. Because the only definitively primary responses are in previously uninfected infants and young children receiving their first vaccination, we have initiated an effort to resolve some of these ambiguities with the pilot study reported here.

Analyses of immune responses in infants require methods that can derive sufficient information from the small blood samples that can be drawn ethically. While previous systems immunology efforts have examined the immune system of infants, for example by use of cytometry by time-of-flight (7), they do not generate information about specificity of the elicited cellular and humoral responses. In this study, we have analyzed cellular and serological influenza antibody repertoires of two infants after their first quadrivalent inactivated influenza vaccine (IIV4) immunization and followed the development of HA-specific immune responses over two years. One of the two infants, who received a prime but not a boost in Year 1, had a documented IBV infection between sample collection in Year 1 and Year 2, providing an indirect comparison of the responses to vaccination and infection. In this infant, there was a shift after the infection from an IAV- to an IBV-dominated Bmem repertoire. The other infant, who received both prime and boost in Year 1, had more balanced responses. In both infants, Year 2 somatic hypermutation (SHM) levels approached those observed in adults. The results suggest that IIV4 may provide a less subtype-dominant, although potentially weaker, imprint than does (monovalent) infection. A more definitive conclusion will require obtaining and analyzing samples from more than the two subjects in this initial pilot study.

## Results

### Study Design

We enrolled two infants -- designated “Infant 1” and “Infant 2” -- with exposure histories shown in Fig. 1A. Both infants received IIV4 concurrently with other vaccines during their first two years (Table S1). Infant 1 received only a prime IIV4 in Year 1 and contracted a confirmed, adventitious IBV infection between Year 1 and Year 2 IIV4. Yamagata lineage IBVs were more prevalent than Victoria lineage IBVs both throughout the influenza season in question (2016-17) and during the week of diagnosis (https://www.cdc.gov/flu/weekly/index.htm). Thus, the infecting virus was likely a Yamagata-lineage IBV, but as both lineages were circulating, we cannot be certain. Infant 2 followed the standard IIV4 vaccination schedule, with a prime and a boost in Year 1. Blood samples were collected at pre- and post-vaccination time points as indicated in Fig. 1A,B.

**Fig. 1.**
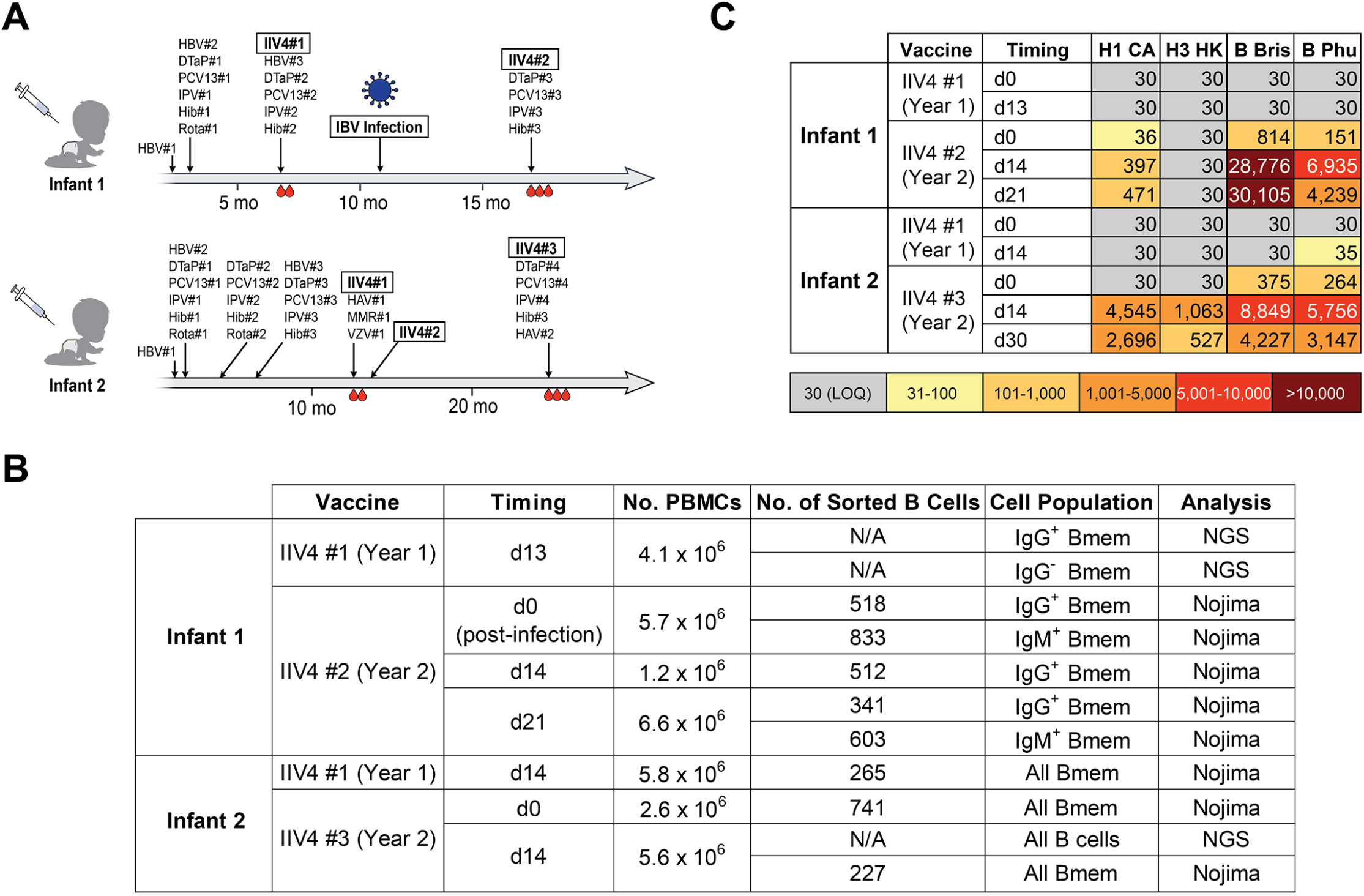
Study design and sample description. **A**. Immunization and influenza infection history for the two infants enrolled in the study. Blood drop symbols indicate the blood draws analyzed in the study. HBV: hepatitis B vaccine; DTaP: diphtheria, tetanus, and pertussis vaccine; PCV: pneumococcal vaccine; IPV: polio vaccine; Hib: haemophilus influenzae type B vaccine; Rota: rotavirus vaccine; IIV4: quadrivalent inactivated influenza vaccine; HAV: hepatitis A virus vaccine; MMR: measles, mumps, and rubella vaccine; VZV: varicella-zoster virus vaccine. **B**. Summary of B cells analyzed in this study. **C**. Plasma IgG binding titers against HAs included in the IIV4. Values indicate ED_50_ from ELISA. H1 CA: A/California/2009 X181; H3 HK: A/Hong Kong/2014 X263B; B Bris: B/Brisbane/60/2008; B Phu: B/Phuket/3073/2013; LOQ = limit of quantitation.

Although the different sampling histories for the two infants studied here diverged in Year 1, with the lack of a booster vaccination for Infant 1, they converged in Year 2, with the “boost” for infant 1 being the intervening IBV infection. Thus, by the time of the Year 2 vaccination, both subjects had two exposures -- in one case, a primary vaccination was followed several months later by infection; in the other, primary vaccination was followed one month later by a second, identical boost vaccination. All the vaccinations were with split vaccines -- *i*.*e*., protein antigens; the infection was an “immunization” with intact virions. The two infants were a year apart in age, and hence in immunization schedule. The 2016-17 and 2017-18 influenza vaccines differed only in their H1N1 components; the 2017-18 and 2018-19 vaccines differed in their H3N2 components and in their Victoria-lineage IBV components.

We measured HA-binding titers of plasma IgG at each time point by ELISA (Fig. 1C). In Year 1, both infants had titers below the limit of quantitation for any of the IIV4 HA components. In Year 2, Infant 1 had a marked increase in IgG titers for IBV HAs and substantially lower titers for IAV HAs (Fig. 1C); Infant 2 had similar titers for both.

We investigated the repertoires of both circulating antibody and HA-specific memory B cells (Bmem). For cellular responses, we profiled the B cells collected from each time point by NextGen sequencing (NGS) and by clonal expansion and sequencing in Nojima cultures (8) (Fig. 1B). To analyze plasma antibody repertoires, we isolated plasma IgG from the Year 2 post-vaccination samples only, as there were insufficient amounts of HA-reactive IgG in plasmas collected in Year 1. We identified the sequences and quantities of each HA-reactive plasma IgG clonotype using the Ig-Seq workflow, as in our previous work (9-12).

### Analysis of the Overall B-Cell Repertoire

Affinity maturation, a consequence of SHM and antigen-driven selection, and antibody class-switching are hallmarks of B cell response to protein antigens. We found no significant increase in the *IGHV* SHM levels for all non–class-switched B cells in the two infants from Year 1 to Year 2 (Fig. 2A). The average SHM levels for non–class-switched B cells at all time points (∼1.5%-2.5%) are similar to those found in adults (13-15), in accord with a published study (16). For all class-switched B cells, the average *IGHV* SHM levels in both infants increased significantly from Year 1 to Year 2 (Fig. 2B), and more prominently in Infant 2 (from 3.1% to 5.6%) than in Infant 1 (from 3.1% to 4.1%). The difference might be due to the greater number of vaccinations (28 for Infant 2; 18 for Infant 1) that Infant 2 received during the time period represented here (Fig. 1A); Infant 2 was also ∼6 months older than Infant 1 at comparable times of blood draw.

**Fig. 2.**
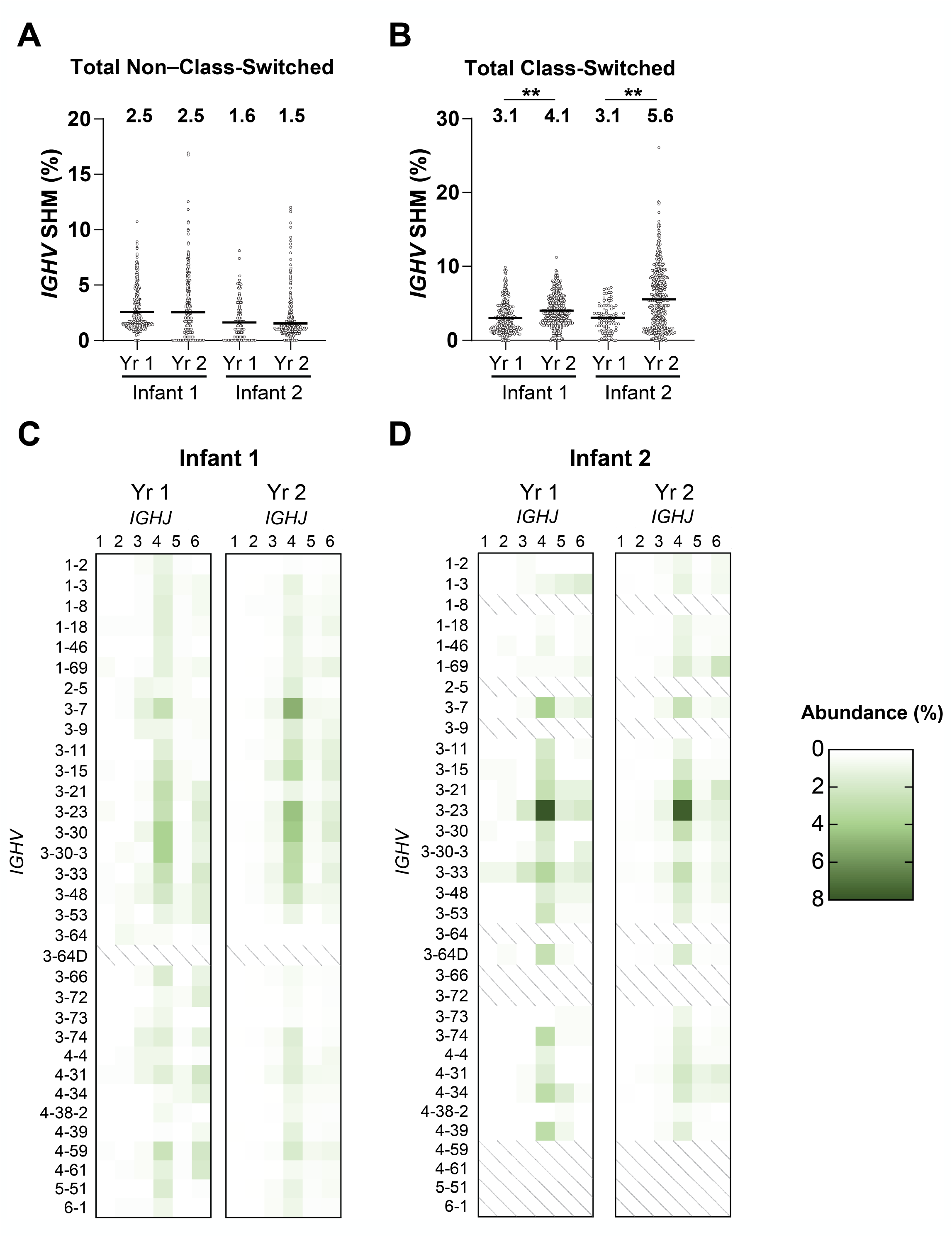
Total B cell repertoire. **A-B**. The *IGHV* SHM levels of non–class-switched (**A**) and class-switched (**B**) total post-vaccination B cells. Median values are represented by horizontal lines and mean values are included above each dataset. Statistical significance was determined by a Mann-Whitney U test (** *P* < 0.01). **C-D**. *IGHV-IGHJ* usage of post-vaccination B cells across two years for Infant 1 (**C**) and Infant 2 (**D**). *IGHV* with >1% abundance in either infant was included in the dataset.

B cell receptors (BCRs) from Bmem cells had similar overall *IGHV* distributions in both children (Fig. 2C,D), with some prevalence of *IGHV 3-7, IGHV 3-23, and IGHV 3-33*; *IGHJ4* was the most abundant J segment. Other molecular features, such as CDRH3 length, hydrophobicity, and charge, which can influence antibody engagement with a variety of different antigens, did not differ significantly between two infants or among time points (Fig. S1) and were similar to the corresponding characteristics observed in adults (17).

### Binding Profiles of HA-specific Memory B cells

We characterized the binding of HA-specific Bmem at each time point by antigen-independent single-cell sorting of Bmem from peripheral blood mononuclear cell (PBMC) samples (Fig. S2) into Nojima-culture wells (8). We then profiled the binding specificities and avidities of the secreted antibodies using a panel of IAV and IBV HAs in a multiplexed bead assay (Supplementary Data 1 and Fig. S3) (18, 19). We determined the HA-specificity and binding avidity by calculating an avidity index (AvIn) for each antibody, as previously described (8). Because Bmem collected from Infant 1 in Year 1 was analyzed only by NGS without the Nojima-culture step and the ensuing multiplexed bead assay, we expressed representative recombinant antibodies selected based on the highest read counts and evaluated their antigen-binding properties by ELISA (Table S2).

We assigned the binding specificities of non–class-switched (Fig. 3A) and class-switched (Fig. 3B) Bmem cells as IAV-specific or IBV-specific based on their relative AvIn values (8). A small population of Bmem cells bound both IAV and IBV HAs with similar, usually low, AvIn, and we designated them as IAV+IBV cross-reactive. The sampled response of Infant 1 shifted from entirely IAV HA-specific (non–class-switched: 3/3; class-switched: 5/5) in Year 1 post-vaccination (after IIV4 #1) to largely IBV HA-specific (non–class-switched: 12/13; class-switched: 6/10) in Year 2 pre-vaccination (before IIV4 #2), the earliest sample time after the IBV infection. The lineage (Yamagata or Victoria) of that infection was not determined. The IBV-biased outcome continued after the Year 2 IIV4 #2 immunization (non–class-switched: 2/2, class-switched: 15/23). We conclude that the IBV infection caused the shift in Bmem reactivity. For Infant 2, binding specificities were more evenly distributed at each sampling time (Fig. 3A,B).

**Fig. 3.**
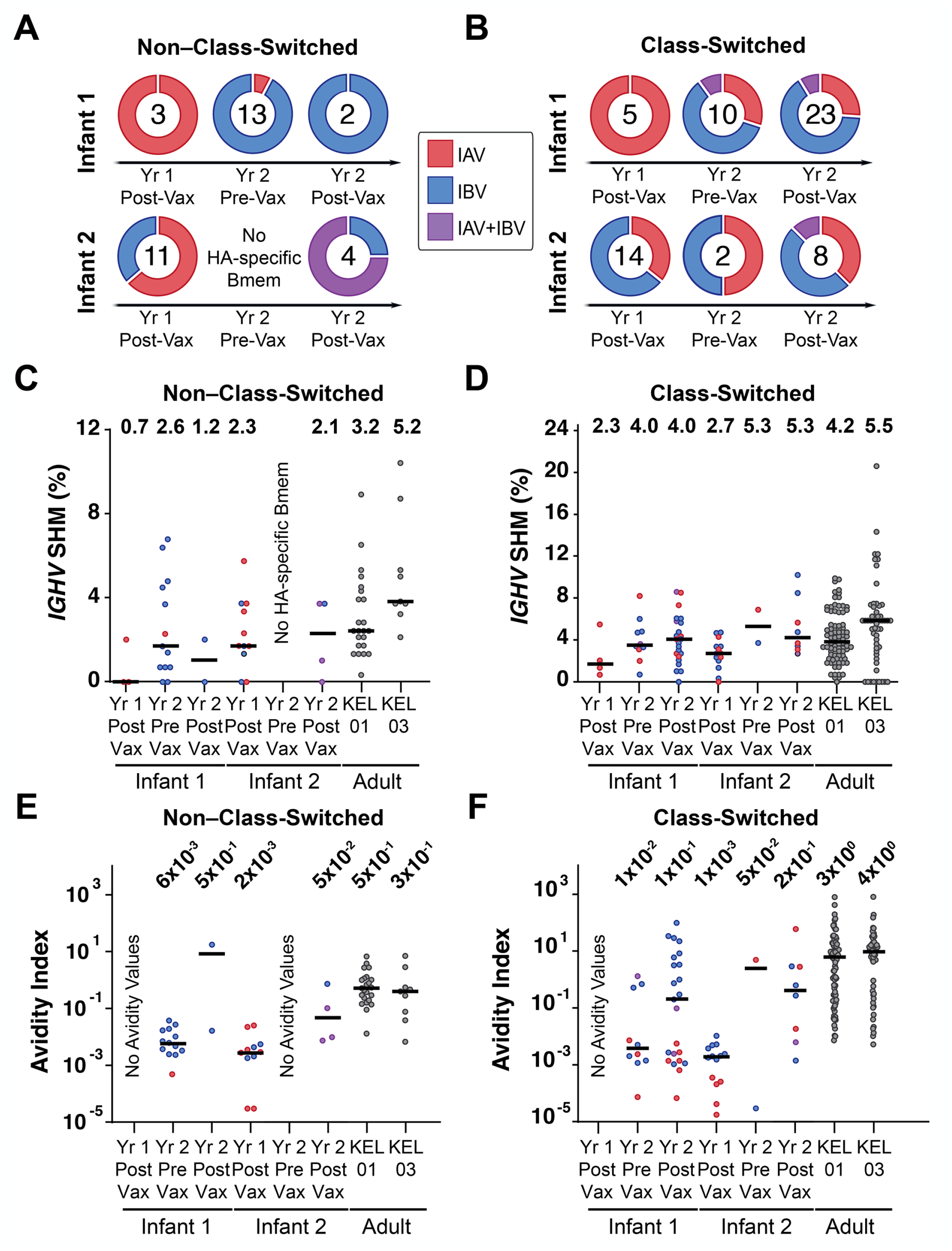
Features of HA-specific Bmem. **A-B**. Binding specificities of HA-specific non–class-switched (**A**) and class-switched (**B**) Bmem. Doughnut plots represent percentage of Bmem that are IAV-specific, IBV-specific, or cross-reactive to both (IAV+IBV). Numbers indicate sample size in each timepoint. **C-D**. *IGHV* SHM levels of HA-specific non–class-switched (**C**) and class-switched (**D**) Bmem. **E-F**. Avidity index of HA-specific non–class-switched (**E**) and class-switched (**F**) Bmem. For **C-F**, median values are represented by horizontal lines and mean (**C-D**), or geometric mean (**E-F**) values are included above each dataset. Two adult subjects studied previously are designated KEL01 (male, age 39) and KEL03 (female, age 39). Red, blue, and purple dots represent IAV-specific, IBV-specific, and IAV+IBV Bmem, respectively.

The mean AvIn of both non–class-switched and class-switched Bmem (Fig. 3E,F) increased across time points, as expected from affinity maturation of HA-specific Bmem. In Infant 1, most of the high-avidity Bmem were IBV-specific, and the (geometric) mean and median AvIn of all the IBV-specific, Year 2 post-vaccination Bmem (∼1×10^0^ and ∼3×10^0^, respectively) in that infant (Supplementary Data 1) were indistinguishable from those of the control adults from two previously studied subjects reporting repeated seasonal influenza vaccination (18) (Fig. 3F). Infant 2 had more evenly distributed binding specificities and avidities after the Year 2 vaccination.

We analyzed post-vaccination SHM levels in both years, for HA-specific Bmem cells. The class-switched, HA-specific Bmem SHM increased from 2.3% and 2.7% in Year 1, for Infant 1 and Infant 2, respectively, to 4.0% and 5.5% in Year 2, with essentially no change in SHM for non-class-switched Bmem cells (0.7% to 1.2% for Infant 1 and 2.3% to 2.1% for Infant 2) (Fig. 3C,D). Adult controls had 3.2% and 5.2% average SHM levels for non–class-switched and 4.2% and 5.5% for class-switched HA-specific Bmem cells. We found no significant correlation between *IGHV* SHM and AvIn (Fig. S4), consistent with previous observations (20, 21).

Because both infants received DTaP vaccinations concurrently with their Year 2 IIV4 immunizations (Fig. 1A), we used tetanus toxoid (TT) as a control in our panel for the multiplexed bead assay and to compare SHM levels in TT-specific Bmem cells as an independent measure of the infants’ response to repeated (2 or 3) immunizations with a single immunogen. Overall frequencies of TT-reactive Bmem cells (Table S3) were similar for the two infants; average SHM levels (*e*.*g*., 4.0% and 7.1% in Infant 1 and Infant 2, respectively for Year 2 post-vaccination; Fig. S4) were similar, in each infant, to HA-specific SHM, even after 4 DTaP doses in the case of Infant 2 (Fig. 1A).

### HA-specific Plasma Antibody Repertoire

Ig-Seq (9-11, 16) profiling of the IAV and IBV HA-reactive plasma IgG repertoires from the Year 2 post-vaccination time point (Supplementary Data 2) showed that the serological antibody repertoires of both infants were oligoclonal, with 8 (Infant 1) and 4 (Infant 2) IAV HA-reactive clonotypes and 41 (Infant 1) and 16 (Infant 2) IBV HA-reactive clonotypes (Fig. 4). With the exception of the IBV HA-specific repertoire of Infant 1, the total number of distinct antibody clonotypes specific for each HA was much smaller than previously observed in adults, which ranged between 40 and 147 at post-vaccination time points (11). In both infants, one highly abundant clonotype dominated the IBV HA-specific repertoire, comprising 42% and 61% of the entire repertoire in infants 1 and -2, respectively; the third most abundant clonotype accounted for only 4.3% and 2.3% of the repertoire (Infant 1 and Infant 2, respectively). The distribution of the IAV HA-specific repertoire was less polarized.

**Fig. 4.**
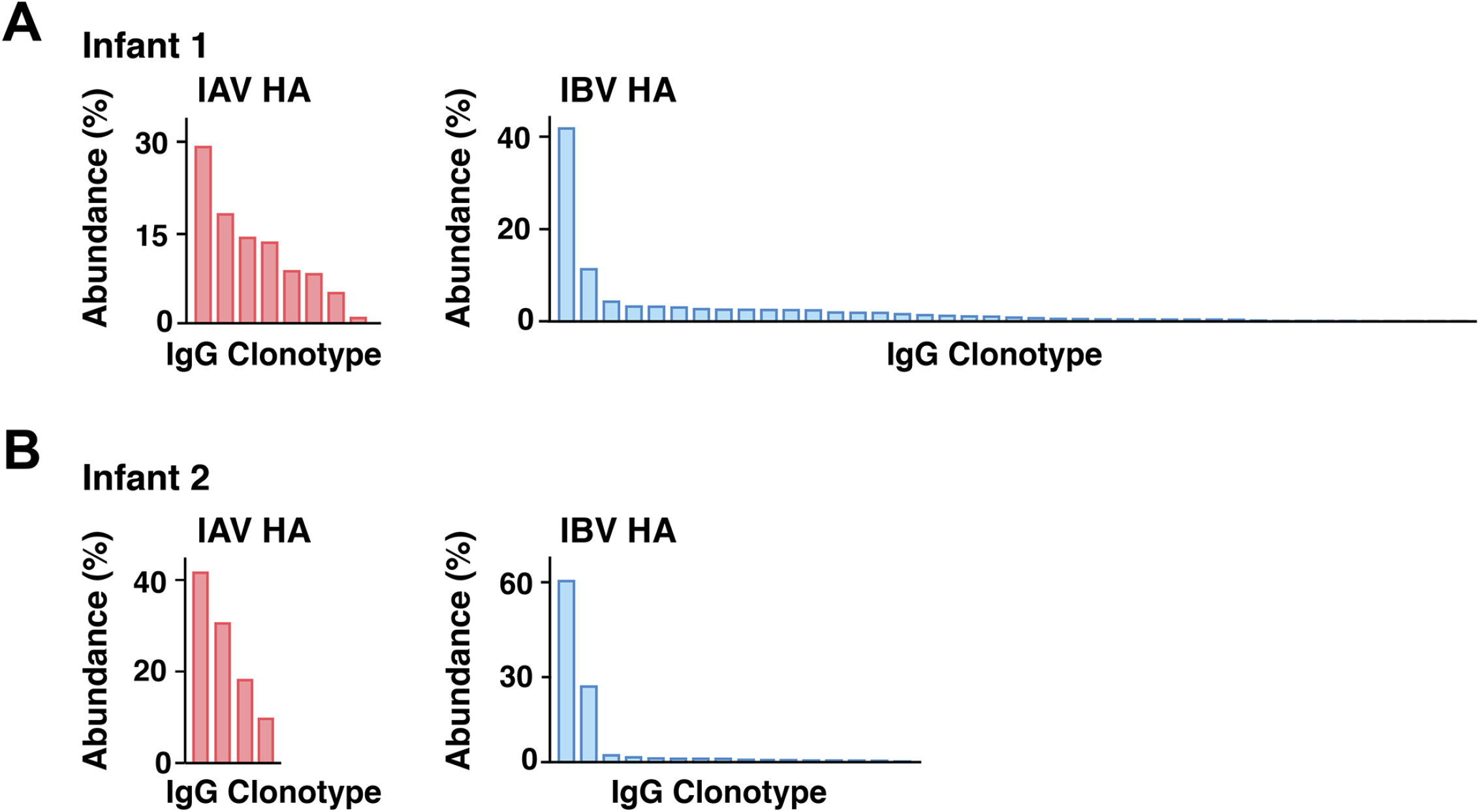
Summary of IAV and IBV HA-reactive plasma IgG repertoire. **A-B**. The composition of IAV and IBV HA-reactive plasma IgG antibody repertoires of Infant 1 (**A**) and Infant 2 (**B**). Each bar represents a unique antibody clonotype, and their heights correspond to the relative abundance of each clonotype in the IAV or IBV HA-reactive IgG repertoire.

The most abundant IBV HA-specific plasma IgG clonotype from Infant 1 is clonally related to a Year 2 post-vaccination Bmem BCR that binds trimeric HA (but not the head alone) with high avidity (AvIn ∼0.2) and broad specificity for B/Phuket/2013, B/Florida/2017, and B/Brisbane/2008 and 3.4% SHM (Supplementary Data 1). The second most abundant clonotype is clonally related to a Year 2 pre-vaccination Bmem BCR with high avidity (AvIn ∼0.7) for B/Phuket/2013. The 3.4% SHM level of this clone, like that of the third most abundant, both presumably affinity matured in response to the IBV infection, was considerably higher than the SHM levels of the most abundant clonotypes in the IAV HA-specific repertoires of both infants 1 and 2 (1.7% and 1.4%) and in the IBV HA-specific repertoire of Infant 2 (0.9%) (Supplementary Data 2).

## Discussion

Technologies such as single-cell NGS and Ig-Seq have enabled detailed molecular dissection of human immune responses to vaccination and infection (11, 16). Before widespread implementation of infant influenza vaccination, most children had experienced an IAV or IBV infection by the age of 3 or 4 (22); previous studies characterizing immune responses in adults to later vaccinations were therefore in the context of pre-existing immune memory. The principal challenge, for extending such studies to true primary responses in infants, is how to extract as much information as possible from the small volumes of blood that can be drawn. Because of restricted blood draw volumes from infants (no more than 0.7 mL/kg for infants and children, compared to 40-50 mL typically drawn from adult subjects), we maximized the depth of BCR repertoire analysis by combining NGS and Nojima culture expansion and sequencing (Fig. 1B). We also performed one affinity purification step using mixed IAV HAs and another using mixed IBV HAs to define IAV and IBV HA-reactive plasma IgG repertoires, respectively. The results presented here illustrate that our approach can lead to meaningful conclusions, despite the limited sampling.

The SHM levels of the overall Bmem repertoire in the infants studied here reflects both affinity maturation in response to the many vaccines in the standard series and environmental exposure to antigens that children commonly encounter. It conforms to patterns previously documented for overall development of B cell repertoire (23-26). For example, a recent examination of the immunological characteristics of childhood B cell development (16) found that by age 3, infants had adult levels of SHM in non–class-switched B cells and levels in class-switched B cells that approached the 7-8% typical of adults (15). Our data show, specifically for HA, that the range of SHM levels in Bmem cells 2-3 weeks after the Year 2 boost was likewise comparable to the range for similar vaccine responses in adults after multiple antigen exposures by vaccination and infection. The specific response to influenza we have analyzed gives a closer view of affinity maturation of HA-reactive antibodies after successive boosts of a particular antigen from vaccination and infection.

For Infant 1, IBV infection shifted Bmem with AvIn above background from primarily IAV HA-reactive in Year 1 to substantially IBV-directed in Year 2 (Fig. 3A,B). The Year 2 vaccination retained and boosted this IBV-biased Bmem repertoire. To the extent that the IBV infection occurred on a nearly IBV-naive background, the single immunization having generated undetectable numbers of Bmem cells, imprinting by infection (IBV) apparently resulted in a type-dominated response, partly eclipsing whatever IAV imprint the Year 1 vaccination had produced. For Infant 2, the Year 2 vaccination probably retained the IAV/IBV balance of the primary response, but the total number of B cells retrieved in the pooled 2- and 3-week post-vaccination samples (Table S3) was too modest to support a strong conclusion.

For both infants, the average SHM levels after the initial, Year 1 vaccination (IIV4 #1), 2.3% and 2.7%, respectively, were similar to those (upper level ca. 2.5%) generally found for primary responses to vaccination with a novel antigen (27-30). IBV infection of Infant 1 may have produced more extended affinity maturation -- a mean of 4% SHM pre-vaccination in Year 2 -- although we cannot rule out that some of the Bmem cells isolated at that time point were descendents of cells elicited by the Year 1 immunization. There is evidence for relatively long-lived GCs (several months) following infection with SARS-CoV-2 (31-34) and correspondingly higher SHM levels at the end of the active response. The class-switched, IBV HA-specific Bmem cells isolated from Infant 1 after the Year 2 immunization had avidities similar to the average avidity of adult influenza HA-specific BCRs, indicating that the Bmem outcome of infection followed by a single vaccination boost is nearly indistinguishable, in SHM and AvIn from the responses of adults imprinted by infection but vaccinated (and perhaps infected) multiple times since then.

Plasma antibodies 2-3 weeks after a boost reflect the plasmablast burst that peaks at about one week, derived from reactivated Bmem differentiating into antibody secreting cells, on the background of contributions of LLPCs from all previous exposures. The plasma antibody repertoire, in the two infants we studied, was more restricted than in adult donors and dominated by an individual clonotype in the IBV-reactive plasma IgG. It remains undetermined how the repertoire develops in subsequent years and in response to subsequent exposures. Studying the course of immunity in a larger number of subjects and over longer times will be necessary to determine how typical are the differences we have found here and, if they are indeed more general, whether subsequent exposure modify their respective characteristics.

## Methods

### Study Approvals and Volunteers

The study procedures, informed consent, and data collection documents were reviewed and approved by the Duke Health IRB (Pro00020561, initial approval 2009). We enrolled two infants who were vaccinated during their first year and boosted a year later (Table S1). From Infant 1, blood was collected on days 0 and 13 following IIV4 #1 in Year 1 and days 0, 14, and 21 following IIV4 #2 in Year 2. From Infant 2, blood was collected on days 0 and 14 following IIV4 #1 in Year 1, and days 0, 14, and 21 following IIV4 #3 in Year 2. Blood samples were processed into PBMC and plasma, then aliquoted and stored at -150°C for future analysis.

### Flow Cytometry

PBMC in RPMI containing 10% FBS were incubated with excess mouse IgG1 (MG1K; Rockland), to block nonspecific binding, and then labeled with fluorochrome-conjugated mAbs. The following human surface antigen specific mAbs, purchased from BD Biosciences, BioLegend or Thermo Scientific were used: anti-human IgM-FITC (MHM-88), CD3-PE-Cy5 (UCHT1), CD14-Tri (TuK4), CD16-PE-Cy5 (3G8), CD19-PE-Cy7 (HIB19), IgG-APC (G18-145), IgD-PE (IA6-2), CD27-BV421 (M-T271), and CD24-BV510 (ML5). Labeled cells were sorted by FACS Aria II with Diva software (BD Biosciences). Flow cytometric data were analyzed with FlowJo software (Treestar Inc.). Total Bmem cells (CD19^+^Dump^−^CD27^+^CD24^+^), IgG^+^, IgG^−^, or IgM^+^ Bmem cells (Figure S2) were identified as previously described (18, 19, 35). Doublets were excluded from cell sorting by combinations of FSC-A versus FSC-H gating. Cells positive for 7-AAD (BD Bioscience) or positive for CD3, CD14, or CD16 expression were also excluded.

### Single-Cell Cultures

Single Bmem cells were expanded in the presence of MS40L-low feeder cells as previously described (18, 19, 36). Single Bmem cells were sorted directly into wells of 96-well plates and co-cultured for 25 days with the MS40L-low feeder cells (37, 38) in the presence of exogenously-added cytokines. Half of the culture media was replaced twice each week with fresh media containing fresh cytokines. On day 25, culture supernatants were harvested for screening the reactivity of clonal IgG antibodies. Expanded clonal B cells were frozen at -80°C for V(D)J sequence analyses.

### BCR Rearrangement Amplification and Analysis

Rearranged V(D)J gene sequences for human B cells from single-cell cultures were obtained by SMART RT-PCR. Total RNA was extracted from selected samples using Quick-RNA 96 kit (Zymo Research). cDNA was synthesized from DNase I-treated RNA using SMARTScribe™ Reverse Transcriptase (Clontech) at 42°C for 90 min followed by 70°C for 15 min with 0.2 μM each of gene-specific reverse primers (hIgG-RV1, hIgKC-RV1, and hIgLC-RV2) and 1 μM of 5’ SMART template-switching oligo (TSO-bioG). Primers used for RT: hIgG-RV1: GGTGTTGCTGGGCTTGTGA; hIgKC-RV1: CCTGTTGAAGCTCTTTGTGAC; hIgLC-RV2: GTGCTCCCTTCATGCGTGA; TSO-bioG: /5Biosg/CCAAGCTGGCTAGCACCATGACAGrGrGrG which contains biotin at its 5’ end (/5Biosg/) and three riboguanosines (rGrGrG) at its 3’ end.

Synthesized cDNA was then subjected to SMART PCR using Herculase II fusion DNA polymerase (Agilent Technologies) with a common forward primer (5Anchor1-FW1: CAAGCTGGCTAGCACCATGA) and gene-specific reverse primers (hIgG-RV2: GGTCACCACGCTGCTGA; hIgKC-RV3: GCTGTAGGTGCTGTCCTTG; hIgLC-RV3: TGTGGGACTTCCACTGCTC for IgG, kappa, and lambda PCR, respectively). PCR conditions were 95°C for 4 min, followed by 30 cycles at 95°C for 30 s, 55°C for 20 s, and 72°C for 30 s. PCR products were submitted to the Duke DNA sequencing facility to obtain DNA sequences. Primers used for sequencing: hIgG-RV4: GGGAAGTAGTCCTTGACCAG; hIgKC-RV4: CTGGGAGTTACCCGATTGGA; hIgLC-RV4: GCTGGCCGCRTACTTGTTG for IgG, kappa, and lambda, respectively.

### V_H_:V_L_-Paired Sequencing

Bmem cells collected 13 days after IIV4 #1 from Infant 1 and B cells collected 14 days after IIV4 #3 from Infant 2 were analyzed by V_H_:V_L_-paired sequencing as previously described (39, 40). Single B cells were encapsulated in emulsion droplets in a custom flow-focusing apparatus. The droplets contained lysis buffer and poly(dT) conjugated magnetic beads to capture mRNA from single B cells. The magnetic beads with mRNA were collected and emulsified to undergo overlap extension RT-PCR, in which V_H_ and V_L_ transcripts were physically linked and amplified by Xenopolymerase (RTX) (41). V_H_:V_L_ amplicons were extracted, amplified by nested PCR, and sequenced using an Illumina MiSeq (2×300 bp paired end reads).

### HA Expression and Purification

rHA head and rHA full-length soluble ectodomain (FLsE) constructs were cloned into pFastBac vector for insect cell expression (Hi5 cells) as previously reported (6, 42-45). All constructs were confirmed by DNA sequencing at the DNA Sequencing Core Facility at Dana Farber Cancer Institute. The HA FLsE constructs used in ELISA assays contained a thrombin or HRV 3C-cleavable C-terminal foldon tag with either a 6xHis or SBP-8xHis tag. All constructs were purified from supernatants by passage over Co-NTA TALON resin (Takara) followed by size exclusion chromatography on a Superdex 200 Increase (GE Healthcare) in 10 mM Tris-HCl, 150 mM NaCl at pH 7.5. Affinity tags were removed using HRV 3C protease (ThermoScientific) and the protein repurified using Co-NTA TALON resin to remove the protease, tag and non-cleaved protein.

### Multiplex Bead Assay

Specificity and avidity of clonal IgG antibodies in culture supernatants were determined in a multiplex bead assay (Luminex Corp.) as described (18, 36) with modifications. Culture supernatants were serially diluted in 1×PBS containing 1% BSA, 0.05% NaN_3_ and 0.05% Tween20 (assay buffer) with 1% milk and incubated for 2 hr with the mixture of antigen-coupled microsphere beads in 96-well filter bottom plates (Millipore). After washing three times with assay buffer, beads were incubated for 1 hr with PE-conjugated mouse anti-human IgG antibodies (JDC-10, Southern Biotech). After three washes, the beads were re-suspended in assay buffer and the plates read on a Bio-Plex 3D Suspension Array System (Bio-Rad). Antigens and controls included BSA, mouse anti-human Igκ (SB81a, Southern Biotech), mouse anti-human Igλ (JDC-12, Southern Biotech), mouse anti-human IgG (Jackson ImmunoResearch), Streptavidin (Invitrogen), Tetanus Toxoid from *Clostridium tetani* (List Biological Laboratories), NP-BSA (coupled in Kelsoe lab), keyhole limpet hemocyanin (KLH, Sigma), ovalbumin (OVA, Siga), Kynureninase (KYNU, Sino Biological), recombinant protective antigen (rPA, BEI Resources), HIV-1 gp140 JR-FL, mutant KYNU (provided by the Duke Human Vaccine Institute (46), Insulin (Sigma), and a panel of recombinant hemagglutinins (full-length constructs of H1/California/2009 X181, H1/Michigan/2015 X-275 (Vaccine Strain), H1/Michigan/2015 (Circulating Strain), H1/Florida/1993, H1/Singapore/2015 IVR 180, H3/Hong Kong/2014 X263B (Vaccine Strain), H3/Hong Kong/2014 (Circulating Strain), B/Phuket/2013, B/Florida/2017, B/Brisbane/2008, and head constructs H3/Hong Kong/2014 X263B, B/Phuket/2013, B/Florida/2017, B/Brisbane/2008). A control mAb (CR9114) was used to check for any biases in the HAs used (Table S4).

### Plasma Antibody Titers

Serological Titers. Binding to recombinant HA proteins was performed as described (47). High-binding 384-well microtiter plates were coated with recombinant HA protein at a final concentration of 2 μg/mL diluted in 0.1 M NaHCO3 and incubated overnight at 4 °C. The plates were washed with 1× phosphate-buffered saline (PBS) containing 0.1% Tween 20 and blocked for 1 h with blocking buffer (40 g whey protein, 150 mL goat plasma, 5 mL Tween 20, 0.5 g NaN3, 40 mL 25× PBS, brought up to 1 L with water). Plates were washed and 10 μL of diluted plasma (starting at 1:30 and serially diluted in block buffer) added directly to each well, followed by incubation for 1.5 h. Plates were washed and then incubated for 1 hr in blocking buffer without NaN_3_ to which of horseradish peroxidase (HRP)-conjugated secondary antibody goat anti-human IgG Fc specific antibody (Jackson Immunoresearch) had been added at the dilution as recommended by the manufacturer. The plates were washed and developed with 3′,5-tetramethylbenzidine substrate (KPL), plates for 15 min. Development was stopped with 1% HCl (Fisher Scientific). Plates were read on a plate reader (Molecular Devices) at 450 nm. The background for each analyte was determined based on nonimmune plasma. Midpoint (ED50) titers were calculated by applying four-parameter logistic regression to the binding data using the *drc* package in R (48).

### Purification of Total IgG from Plasma and Subsequent Digestion into F(ab’)2

Plasma samples collected following IIV4 #2 (day 14: 130 μL; day 21: 500 μL) from Infant 1 and following IIV4 #3 (day 14: 250 μL; day 21: 500 μL) from Infant 2 were passed through a 1.5 mL Protein G agarose (Pierce) affinity column three times in gravity mode. The column was washed with 20 column volume (cv) DPBS and eluted with 10 mL of 100 mM Glycine-HCl (pH 2.7) and eluate was immediately neutralized with 1.5 mL of 1 M Tris-HCl (pH 8.0). Purified IgG was digested into F(ab’)2 with 200 μg of IdeS per 10 mg of IgG for 6 hr on a thermal mixer (Eppendorf ThermoMixer C) at 37°C. The digest was rotated with 200 μL of Strep-Tactin Superflow resin (IBA Lifesciences) at room temperature (RT) for 1 hr and separated using a Pierce spin column, collecting the F(ab’)2 in the flow-through.

### Antigen-Enrichment of F(ab’)2 and Mass Spectrometry Sample Preparation

rHAs were immobilized on separate columns for the antigen-specific F(ab’)2 enrichment as previously described (11). Briefly, rHAs were separately immobilized on N-hydroxysuccinimide (NHS)-activated agarose resins (Pierce) by overnight rotation at 4°C; for Infant 1, 0.5 mg of mixed H1/Michigan/2015 X-275 and H3/Hong Kong/2014 X263B or mixed B/Brisbane/2008 and B/Phuket/2013 (Vaccine Strains), and for Infant 2, 0.5 mg of mixed H1/Michigan/2015 X-275 and A/H3/Singapore/2016 or B/Maryland/2016 and B/Phuket/3073/2013 (Vaccine Strains) was used to generate two affinity resins per infant. The coupled agarose resins were washed with DPBS, and unreacted NHS groups were blocked with 1M Tris, pH 8 for 30 min at RT. The resin was further washed with 5 mL of DPBS and added to F(ab′)2 sample. Resin and F(ab’)2 mixtures were rotated for 1 hr at RT. Flow-through samples were collected, and the columns were washed with 10 column volumes of DPBS. Antigen-enriched F(ab’)2 was eluted with 1% formic acid in 0.50 mL. Flow-through and elution fractions were assayed by indirect ELISA with the donor-specific rHAs and the depletion of ELISA signal in each flow-through sample evaluated. Elution fractions showing >0.1 OD_450_ were pooled and concentrated under vacuum to a volume of <10 μL, resuspended in 40 μL of PBS, and neutralized with 1M NaOH.

For each enrichment, elution and flow-through samples were denatured in 50 μL of 2,2,2-trifluoroethanol (TFE), and 5 μL of 100 mM dithiothreitol (DTT) at 55°C for 45 mins, then alkylated by incubation with 3 μL of 550 mM iodoacetamide (Sigma) for 30 mins at RT in the dark. Samples were diluted 10-fold with 40 mM Tris (pH 8) and digested with trypsin (1:30 trypsin/protein) for 12 hr at 37°C. Formic acid was added to 1% (v/v) to quench the digestion, and the sample volume was reduced to ∼100 μL under vacuum. Peptides were then bound to a C18 Hypersep SpinTip (Thermo Fisher Scientific), washed three times with 0.1% formic acid, and eluted with 60% acetonitrile, 0.1% formic acid. C18 eluate was dried under vacuum centrifugation and resuspended in 50 μL in 5% acetonitrile, 0.1% formic acid.

### LC-MS/MS

Samples were analyzed by liquid chromatography-tandem mass spectrometry on either a Dionex Ultimate 3000 RSLCnano uHPLC system (Thermo Fisher Scientific) or Easy-nLC 1200 (Thermo Fisher Scientific) coupled to an Orbitrap Fusion Tribrid or Velos Pro mass spectrometer (Thermo Fisher Scientific). Peptides were first loaded onto either Acclaim PepMap RSLC NanoTrap column (Dionex; Thermo Fisher Scientific) or a prior to separation on a 75 μm×15 cm Acclaim PepMap RSLC C18 column (Dionex; Thermo Fisher Scientific) using either a 5–40% (v/v) or 5-32% (v/v) acetonitrile gradient over 95 or 120 min at 300 nL/min. Eluting peptides were injected directly into the mass spectrometer using a nano-electrospray source. The instruments were operated in a data-dependent mode with parent ion scans (MS1) collected at 120,000 or 60,000 resolution. Monoisotopic precursor selection and charge state screening were enabled. Ions with charge ≥+2 were selected for collision-induced dissociation fragmentation spectrum acquisition (MS2) in the ion trap. Sample were run three times to generate technical replicate datasets.

### MS/MS Data Analysis

Donor-specific protein sequence databases were constructed using the donor’s corresponding V_H_ sequences with ≥2 reads, concatenated to a database of background proteins comprising non-donor-derived V_L_ sequences (HD1) (9), a consensus human protein database (Ensembl 73, longest sequence per gene) and a list of common protein contaminants (MaxQuant). Donor-specific spectra were searched against the corresponding donor-specific databases using SEQUEST HT (Proteome Discoverer 2.4; Thermo Fisher Scientific). Searches considered fully tryptic digested peptides only, allowing up to two missed cleavages. A precursor mass tolerance of 5 ppm and a fragment-mass tolerance of 0.5 Da were used. Modifications of carbamidomethyl cysteine (static) and oxidized methionine (dynamic) were selected. High-confidence peptide-spectrum matches (PSMs) were filtered at a false discovery rate of <1% as calculated by Percolator (q-value < 0.01, Proteome Discoverer 2.4; Thermo Fisher Scientific).

Iso/Leu sequence variants were collapsed into single peptide groups. For each scan, PSMs were ranked first by posterior error probability (PEP), then q-value, and finally based on XCorr. Only unambiguous top-ranked PSMs were kept; scans with multiple top-ranked PSMs (equivalent PEP, q-value, and XCorr) were designated ambiguous identifications and removed. The average mass deviation (AMD) for each peptide was calculated as described (10) using data from all injections and the peptides with an AMD >1.7 ppm were removed.

Peptide abundance was calculated from the extracted-ion chromatogram (XIC) peak area as described (9), using peak area values generated by the Precursor Ions Area Detector node in Proteome Discoverer. For each peptide, a total XIC area was calculated as the sum of all unique XIC areas of associated precursor ions, and the average XIC area across replicate injections was calculated for each sample. Peptide PSM was calculated as the sum of the aforementioned PSMs. For each antigen data set, the eluate and flow-through abundances were compared and peptides with ≥5-fold higher signal in the elution sample were considered to be antigen specific.

### Clonotype Indexing and Peptide-to-Clonotype Mapping

V_H_ sequences were grouped into clonotypes based on single-linkage hierarchical clustering as described (9, 11). Cluster membership required ≥90% identity across the CDRH3 amino sequence as measured by Levenshtein distance. High-confidence peptides identified by MS/MS analysis were mapped to clonotype clusters, and peptides uniquely mapping to a single clonotype were considered “informative”. Abundance of each antibody clonotype was calculated by summing the XIC areas of the informative peptides mapping to ≥4 amino acids of the CDRH3 region (*i*.*e*., CDRH3 peptides). For each antibody clonotype, the most abundantly detected CDRH3 peptide (in case where peptides from multiple somatic variants belonging to the same clonotype were detected) was used as a representative CDRH3 sequence.

Clonotype abundance was calculated as the sum of the peptide abundances belonging to a clonotype. When multiple non-overlapping tryptic peptides were available for a clonotype, one peptide was selected and used for the analysis to prevent double-counting and both the PSMs and XICs were summed across collapsed peptides.

### Recombinant Antibody Synthesis, Expression and Purification

Sequences of variable heavy- and light-chain regions for chosen antibodies were codon optimized and synthesized as eBlocks (Integrated DNA Technologies). The synthetic genes were cloned into a pcDNA3.4 vector (Invitrogen) containing human IgG1 heavy and kappa or lambda light chain constant regions, respectively (16, 49). Heavy and light chain plasmids for each monoclonal antibody were co-transfected into Expi293F cells (Thermo Fisher Scientific). After incubating for 5 days at 37°C with 8% CO_2_, the supernatant containing secreted antibodies was centrifuged at 4000×g for 15 min at 4°C, filtered through 0.45 μm syringe filters (Sartorius), and passed three times over a column with 1 mL Protein G agarose resin (Pierce). After washing the column with 20 column volumes of PBS, antibodies were eluted with 10 mL 100 mM Glycine-HCl (pH 2.7) and immediately neutralized with 1.5 mL of 1 M Tris-HCl (pH 8.0). Antibodies were buffer exchanged into DPBS utilizing Amicon Ultra-30 centrifugal spin columns (Millipore).

### Enzyme Linked Immunosorbent Assay (ELISA)

96 well EIA/RIA Assay Microplates (Corning) were coated with 4 mg/mL of antigen at 4°C overnight. Plates were washed with PBST, blocked with 2% BSA in PBS for 2 hr at RT, and washed again. Serially diluted recombinant antibodies or plasmas were added to the plates and incubated for 1 hr. The plates were washed, and 1:5,000 diluted goat anti-human-IgG Fc horseradish peroxidase (HRP) conjugated secondary antibodies (Invitrogen) were added. After a 1 hr incubation, plates were washed and 100 mL TMB-ultra substrate (Thermo Fisher Scientific) added to each well to develop; 50 mL 2M H_2_SO_4_ was added to quench the reaction. Absorbance was measured at 450 nm (OD450) with a SpectraMax i3x plate reader. Data was analyzed and fitted for EC_50_ if necessary, using a four-parameter logistic nonlinear regression model in the GraphPad Prism (9.0) software.

### Statistical Analysis

Statistical analyses were performed using GraphPad Prism 9.0 (GraphPad Software, Inc., San Diego, CA). All statistical tests performed are described in the figure legends, and significance was determined by P≤0.05.

### Data Availability

The nucleotide sequences for heavy- and light-chain variable domains of the antibodies listed in Supplemental Data Set 1 have been deposited in GenBank, with accession numbers OP419769-OP419958. All the Ig-Seq data (raw and processed proteomics datasets along with the reference sequence databases) from this study have been uploaded to MassIVE and can be accessed under the identifier MSV000090057.

## Acknowledgments

The research was supported by NIAID Grant P01 AI089618. We thank all members of the consortium participating in this Program Project for discussions throughout the course of the research reported here. We also acknowledge support from NIH Grant P20 GM113132 (to JL), NSF Graduate Research Fellowship 1840344 (to NCC), and NIAID CIVIC contract 75N93019C00051 (to GB, GG, GCI). SCH is an Investigator in the Howard Hughes Medical Institute.

## Supplementary table, figure, and data-set legends

**Table S1.**
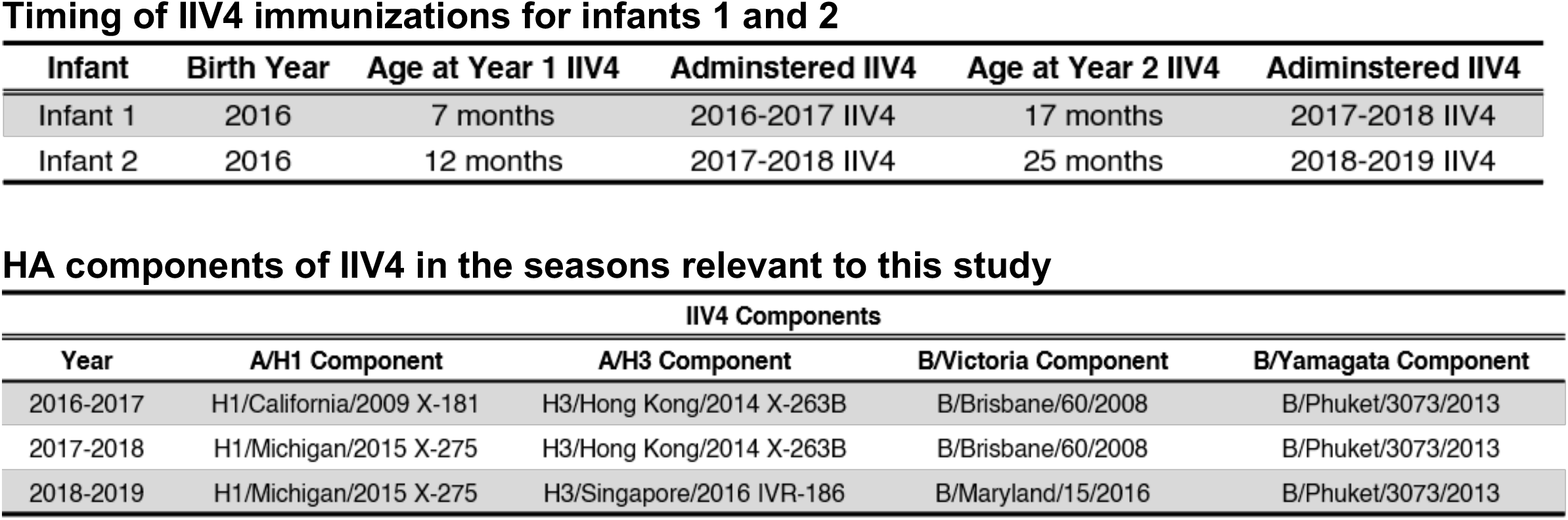
IIV immunizations and their components.

**Table S2.**
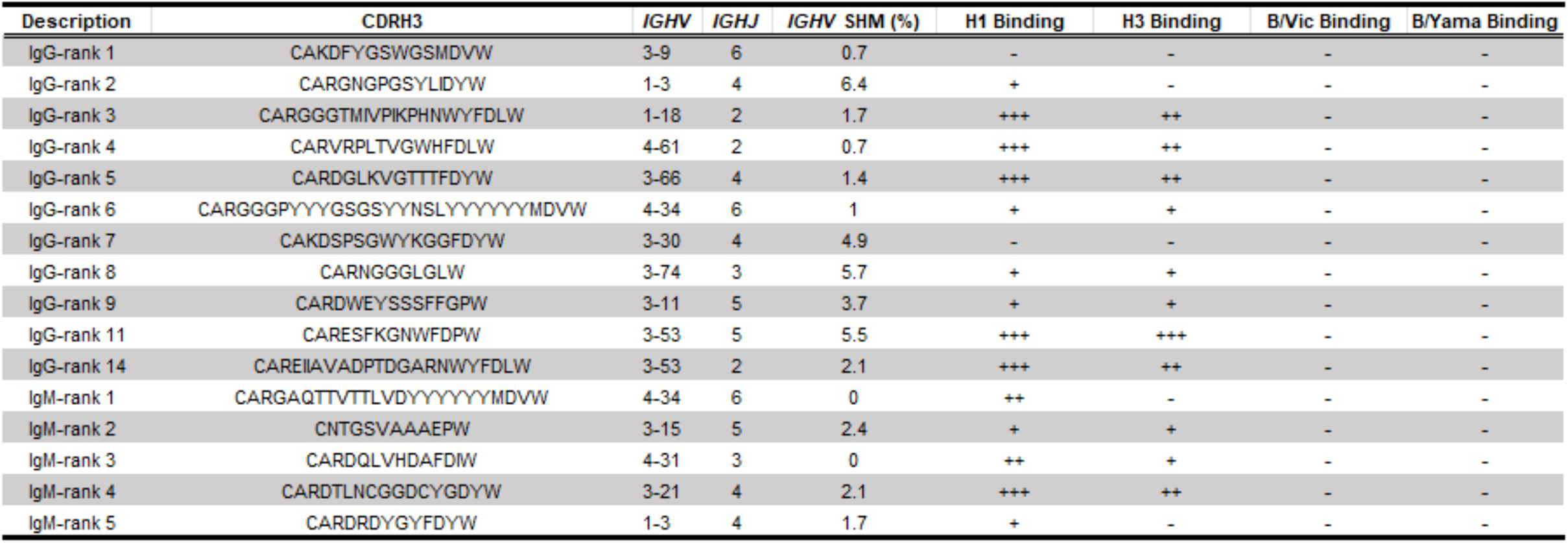
Recombinant mAbs from Infant 1 Year 1 post-vaccination Bmem. Clonotypes with high read counts from the NGS data were made as mAbs. For each selected clonotype, the CDRH3 sequence, *IGHV, IGHJ*, and SHM levels are shown. mAbs were characterized by ELISA to determine their EC_50_ values against HAs. +++: <100 nM; ++: 100-1000 nM; +: detected signal at 1000 nM; -: no signal at 1000nM; H1: A/California/2009 X181; H3: A/Hong Kong/2014 X263B; B/Vic: B/Brisbane/60/2008; B/Yama: B/Phuket/3073/2013.

**Table S3:**
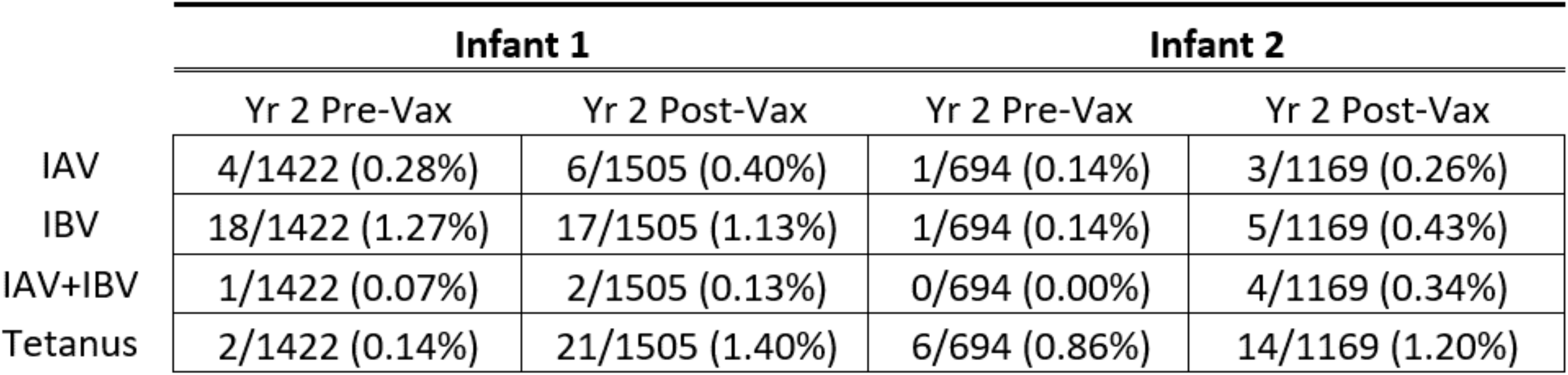
Frequency of HA- and TT-reactive Bmem.

**Table S4:**
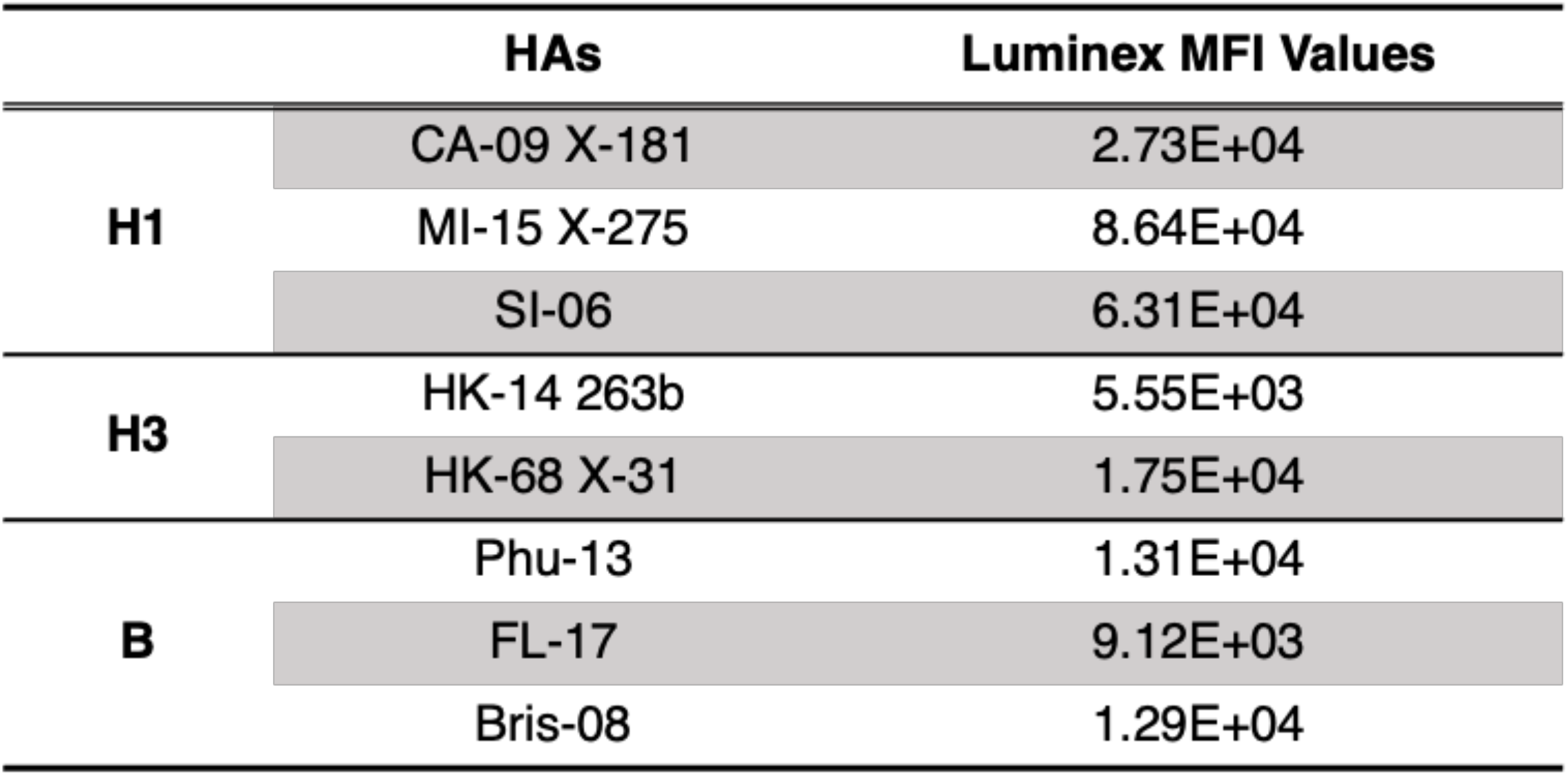
MFI values for CR9114 binding to each HA used in Luminex assay.

**Fig. S1.**
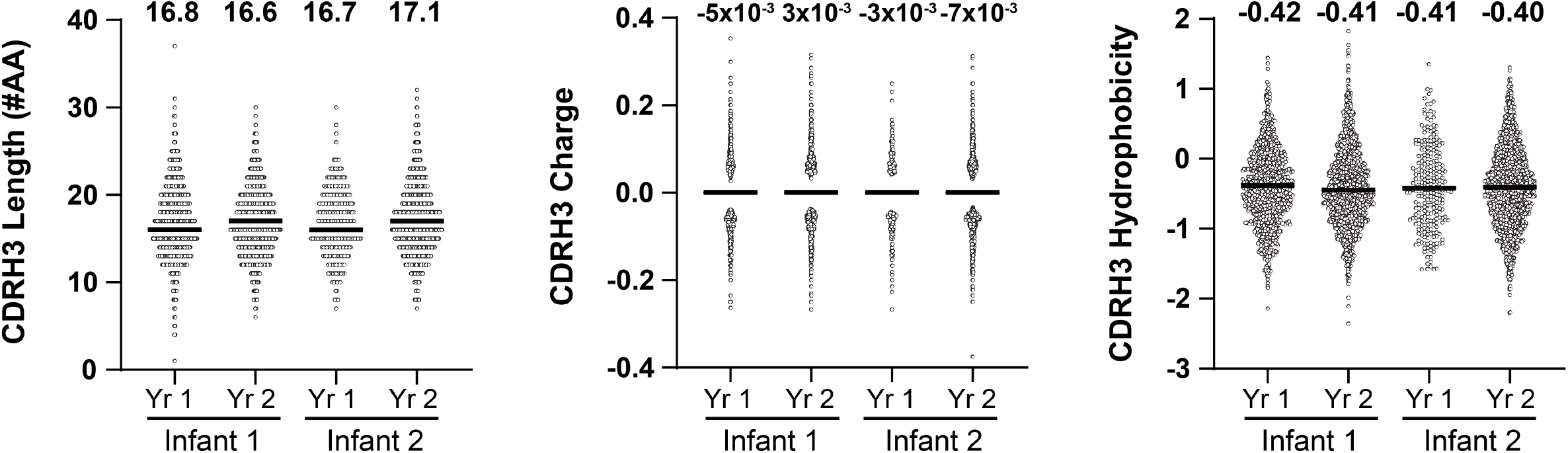
CDRH3 length, charge, and hydrophobicity of total post-vaccination B cells. Median values are represented by horizontal lines and mean values are included above each dataset.

**Fig. S2.**
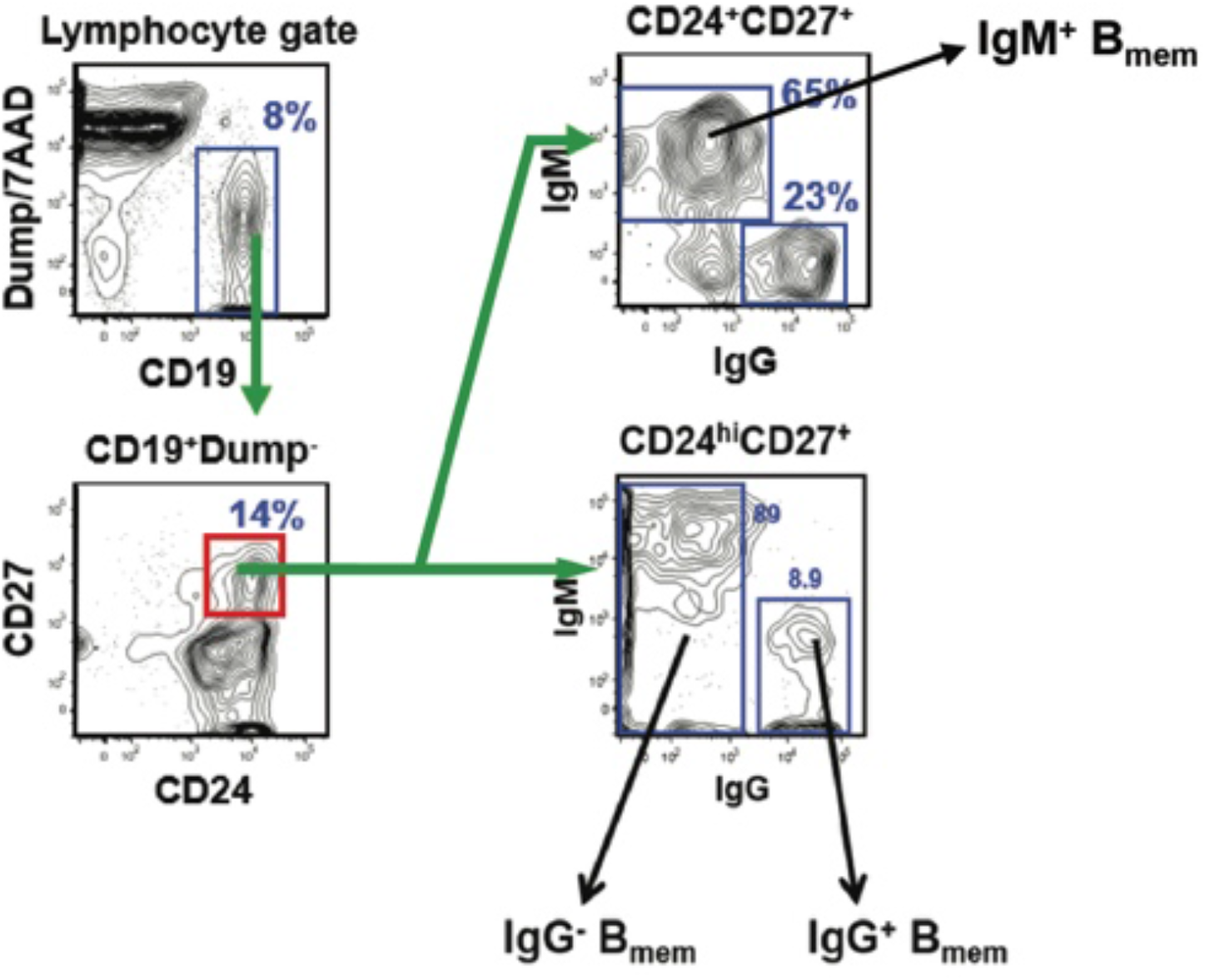
Representative sorting strategy to isolate Bmem for Nojima culturing.

**Fig. S3.**
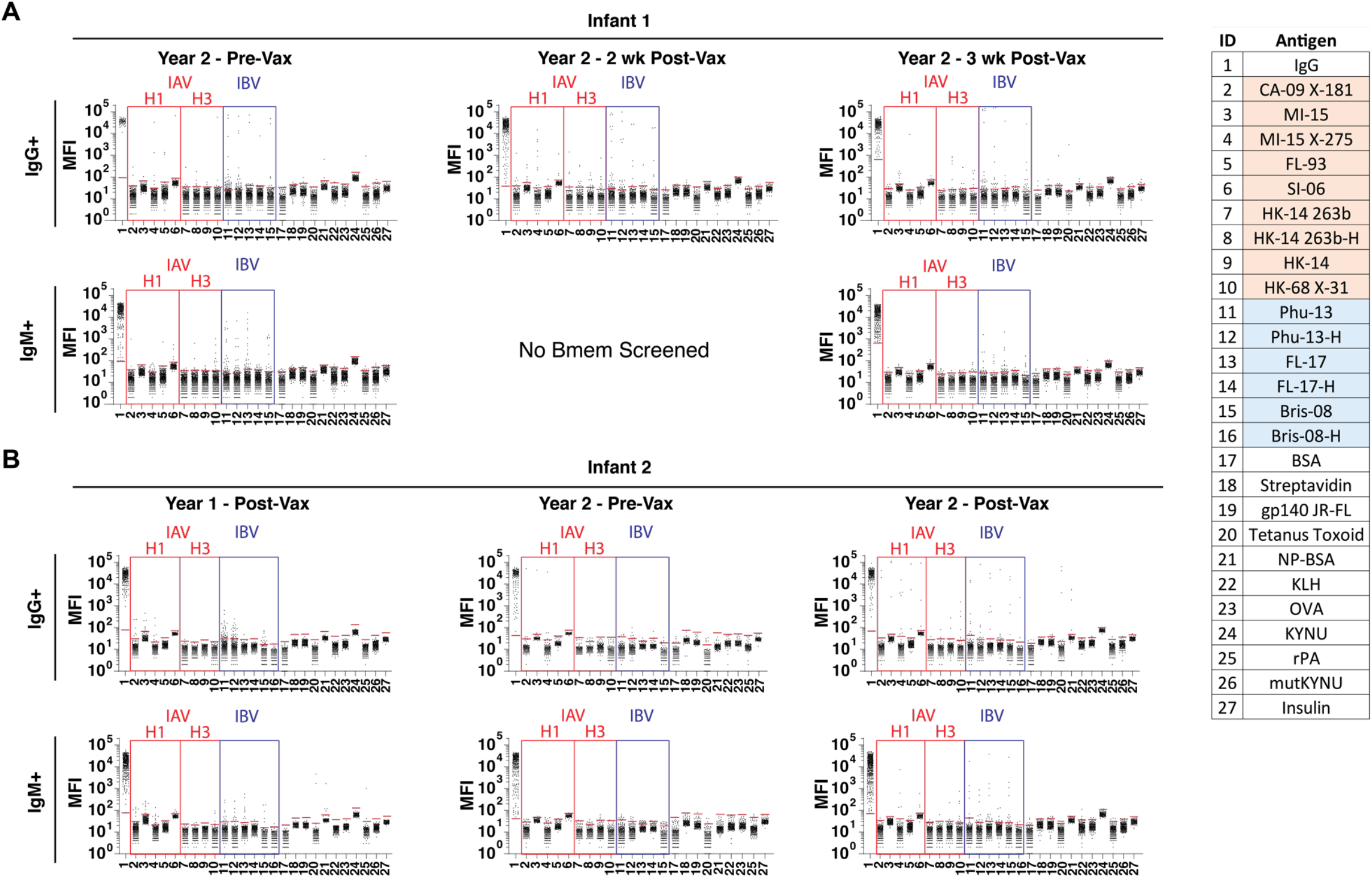
Luminex reactivity assays. Red lines indicate background measurements for each antigen. List on the right shows antigen corresponding to each numbered column. See Materials and Methods (Multiplex Bead Assay) for full name of each HA and control antigen. IgG: mouse anti-human IgG. Antigens marked H are trimeric, head-only constructs.

**Fig. S4.**
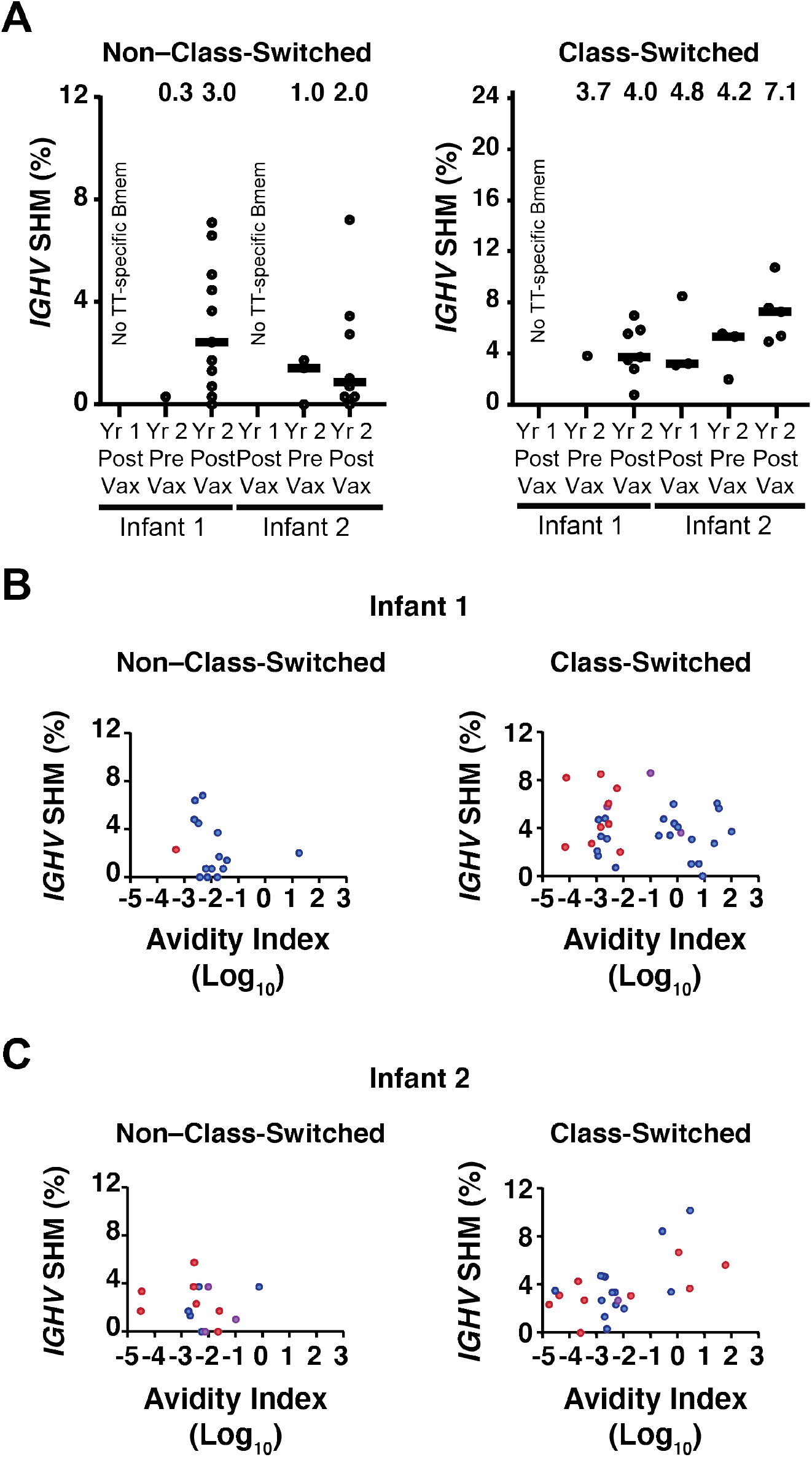
SHM for non-class-switched and class-switched Bmem. **(A) *IGHV* SHM of tetanus-toxin (TT) reactive Bmem**. Median values are represented by horizontal lines and mean values are included above each dataset for TT-specific non–class-switched and class-switched Bmem. **(B and C) Avidity index vs. SHM level of HA-specific Bmem**. Correlation plots for Infant 1 (**B**) and Infant 2 (**C**) are shown for HA-specific non–class-switched and class-switched Bmem. Red, blue, and purple dots represent IAV-specific, IBV-specific, and IAV+IBV Bmem, respectively. Spearman correlation was non-significant (*P* > 0.05) among all subsets.

**Suppl. Data Set 1.**
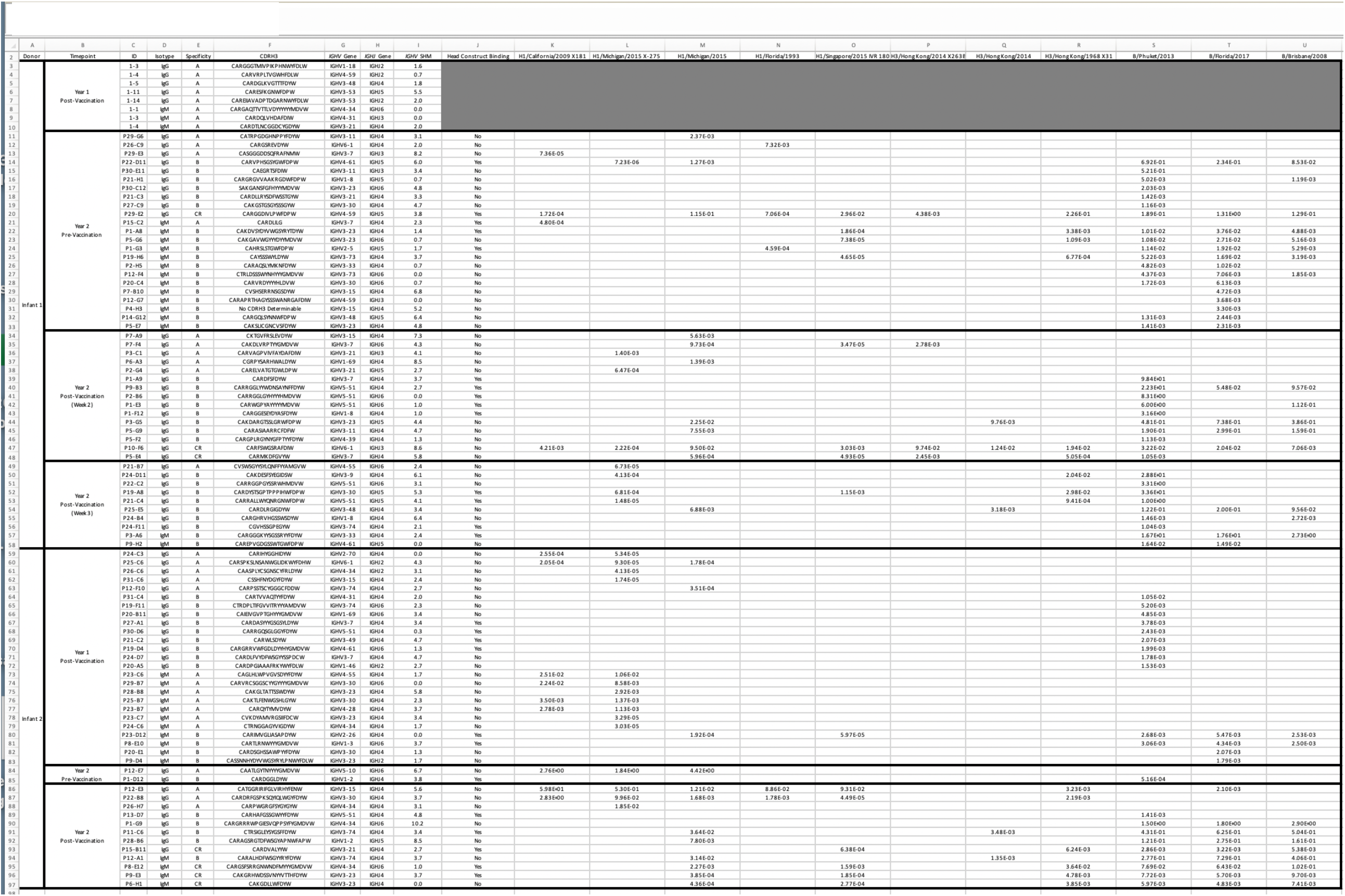
Sequences and molecular features of individual HA-reactive Bmem cells and corresponding Luminex reactivity data. “Head construct” column: “Yes” if the IgG bound *any* of the head-only constructs in the Luminex panel. For full sequences, see Data Availability

**Suppl. Data Set 2.**
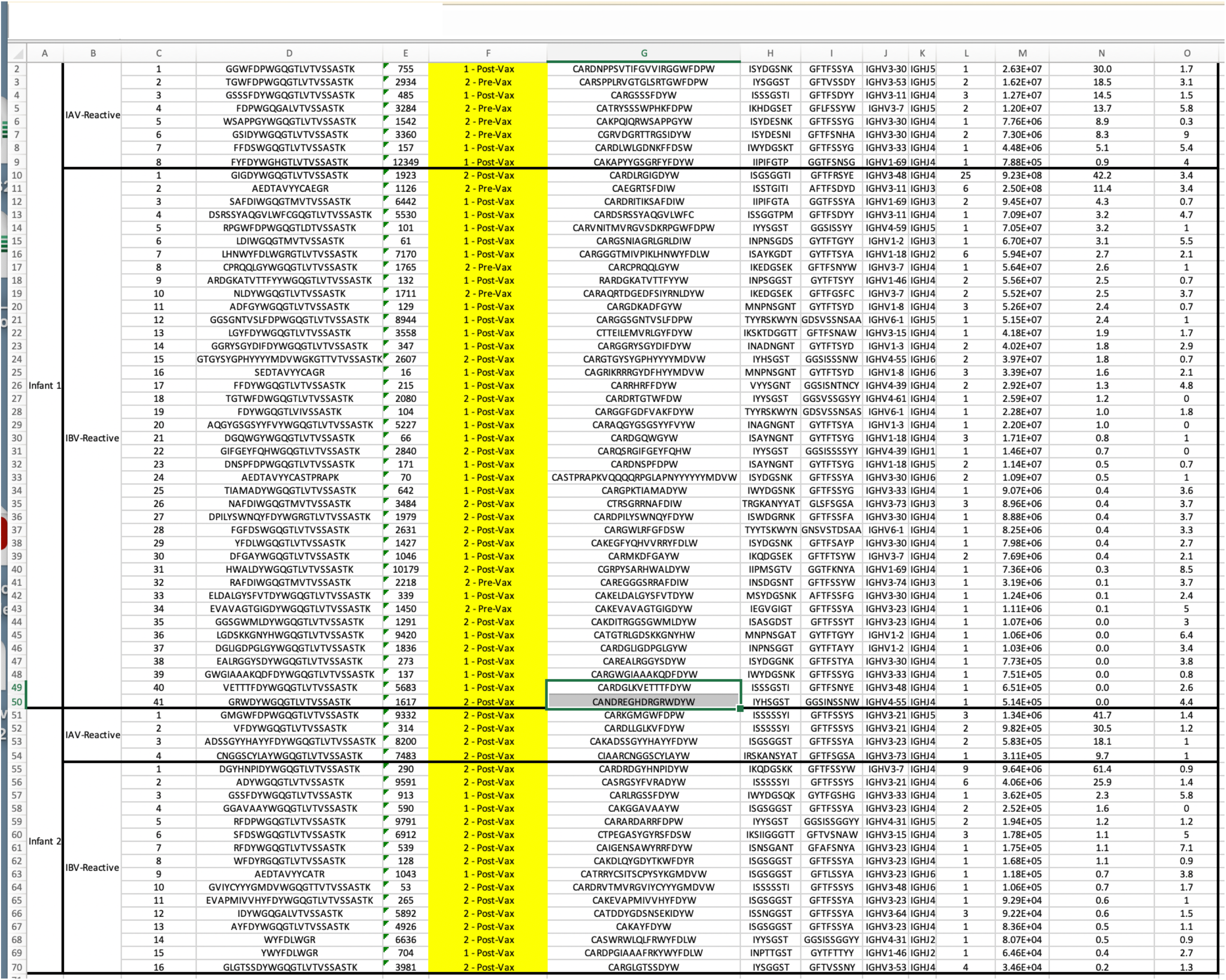
Identities, relative abundances, and molecular features of individual antibody clonotypes in the serological repertoire of IAV- and IBV-reactive antibodies from each infant, as analyzed by Ig-Seq.

## Notes

https://www.ncbi.nlm.nih.gov/genbank/

https://massive.ucsd.edu/

### Competing Interest Statement

The authors have declared no competing interest.

